# Discrete anatomical coordinates for speech production and synthesis

**DOI:** 10.1101/148007

**Authors:** M Florencia Assaneo, Daniela Ramirez Butavand, Marcos A Trevisan, Gabriel B Mindlin

## Abstract

The sounds of all languages are described by a finite set of symbols, which are extracted from the continuum of sounds produced by the vocal organ. How the discrete phonemic identity is encoded in the continuous movements producing speech remains an open question for the experimental phonology. In this work, this question is assessed by using Hall-effect transducers and magnets -mounted on the tongue, lips and jaw- to track the kinematics of the oral tract during the vocalization of *vowel-consonant-vowel* structures. Using a *threshold strategy*, the time traces of the transducers were converted into discrete motor coordinates unambiguously associated with the vocalized phonemes. Furthermore, the signals of the transducers combined with the discretization strategy were used to drive a low-dimensional vocal model capable of synthesizing intelligible speech. The current work not only addressed a relevant inquiry of the biology of language, but also shows the performance of the experimental technique to monitor the displacement of the main articulators of the vocal tract while speaking. This novel electronic device represents an economic and portable option to the standard system used to study the vocal tract movements.

## INTRODUCTION

Among all species humans are the only ones capable of generating speech. This complex process, that distinguishes us from other species, emerges as an interaction between the brain activity and the physical properties of the vocal system. This interaction implies a precise control of a set of articulators (lips, tongue and jaw) to produce a continuous change in the shape of the upper vocal tract^1^. The output of this process is the speech wave sound, which could be discretized and represented by a finite set of symbols: the phonemes. Moreover, the phonemes across languages could be hierarchically organized in terms of articulatory features, as described by the International Phonetic Alphabet^2^ (*IPA*). On the other side of the process, at brain level, intracranial recordings registered during speech production showed that motor areas encode the same set of articulatory features^3^. Then, one missing piece of the puzzle is: how does the continuous vocal tract movement generating speech encode the discrete information?

During the speech production process the displacement of the articulators modifies the vocal tract configuration allowing: *(i)* to apply different filters on the sound initiated by the oscillations of the vocal folds at the larynx (i.e. vowels) and *(ii)* to produce a turbulent sound source by occluding (i.e. stop consonants) or constricting (i.e. fricatives) the tract^4^. Previous works developed biophysical models for this process^5,6^ and tested its capabilities to synthesize realistic voice^7^. In principle, those models could have a high dimensionality, especially due to the many degrees of freedom of the tongue^5^. However, its dimension ranges between 7 and 3, suggesting that a small number of measurements of the vocal tract movements should be able to successfully decode speech and to feed the synthesizers.

In this study the oral dynamics is monitored using sets of Hall-effect transducers and magnets mounted on the tongue, lips and jaw during the utterance of a corpus of syllables (including all the Spanish vowels and voiced stop consonants). By applying a *threshold strategy* on the signals recorded by three sensors it was possible to decode the uttered phonemes well above chance. Moreover, the signals are used to drive an articulatory synthesizer producing intelligible speech. The results disclose that continuous measurements from the oral movements could be represented in a discrete motor-coordinates space; explicitly showing that all steps comprising the speech process can be described in terms of discrete units.

From a technical point of view, the present work represents a benchmark on the state of the art of the measurement techniques used in the speech production field. During the last decades no many improvements have been achieved on the experimental methods used to measure the vocal tract movements. The widely used technique on the field is the electromagnetic articulography^8,9^ (EMA). This equipment produces very accurate measurements but it presents two main disadvantages: it is non-portable and is expensive. The device described in the current work (which is shown to be capable of tracking the vocal tract during continuous speech) represents an alternative method without the problems described above.

## RESULTS

Following the procedure described by Assaneo et al.^10^ sets of Hall-effect transducers and magnets (Figure 1a) were mounted on the upper vocal tract to record the displacement of the articulators (jaw, tongue and lips). More specifically, 3 transducers and 4 magnets were placed on the oral cavity of the participant following the configuration displayed in Figure 1b. The position of the elements was chose in a way that each transducer signal is modulated by a subset of magnets (color code in Figure 1b, see Methods for a more detail). The upper teeth transducer signal represent an indirect measurement of aperture of the jaw, the lips transducer signal represents the roundness and closure of the lips and the palate transducer gives an indirect measure of the position of the tongue within the oral cavity. To diminish the body surface in contact with the glue participants wore plastic molds in their upper and lower dentures (Figure 1a). Just three elements were glued directly on the participants skin: the ones on the tongue and lips. Also, this strategy diminishes the variance on the device configuration between different sessions (the elements on the mold stayed fixed).

**Figure 1.**
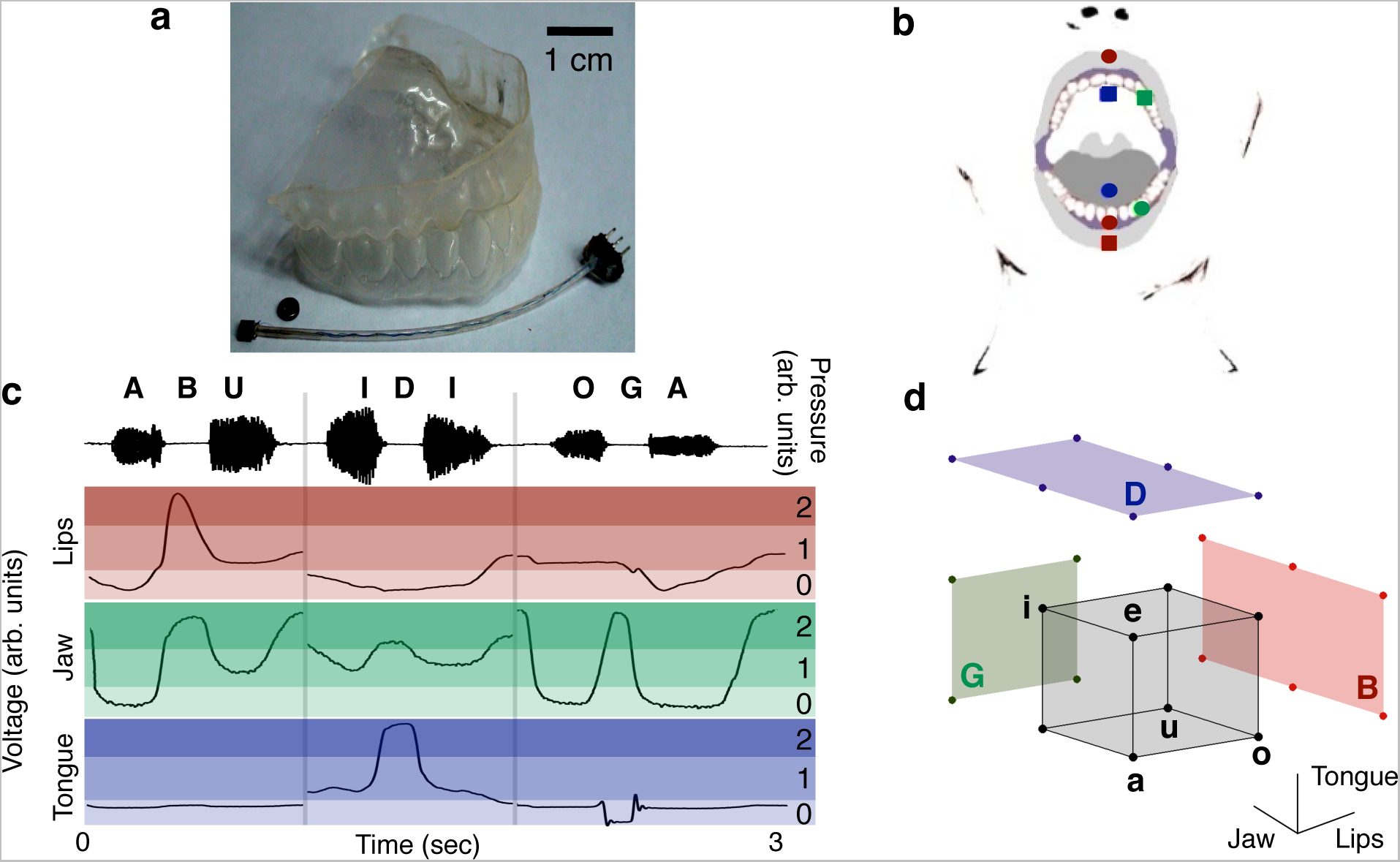
Discrete coordinates are extracted from the time traces of three detectors in the oral cavity. A) The dental plastic replica with a magnet and a Hall-effect transducer. B) Sketch of the position of the set of transducers (squares) and magnets (circles) in the oral cavity. In green the set tracking the jaw -the magnet and transducer were glued to the molds at the upper and lower left canines respectively. In blue the set tracking the tongue -the transducer fixed to the mold at the palate and the magnet to the tip of the tongue. In red the set tracking the lips –the transducer fixed in the center of lower lip, one magnet glued to the lower mold, between the lower central incisors, and other in the center of the upper lip. C) Discretization of the signals. Upper panel: *VCV* phonemes and sound wave. Lower panel: time traces of the transducers signals during the vocalizations. The intensity of the shaded regions differentiates the discrete states. D) States of vowels and consonants in the discrete motor space. The corners of the grey cube represent the vowels and the colored walls de different consonants.

Four native Spanish speakers were instructed to vocalize a corpus of syllables while wearing the device. The transducer signals (*h_J_*(*t*), *h_T_*(*t*) and *h_L_*(*t*) for the jaw, tongue and lips respectively) were recorded simultaneously with the produced speech, an example of the four signals is shown in Figure 1c (see Methods for a complete description of the data acquisition and the preprocessing of the signals). The syllable corpus consists in the 75 possible *vowel-consonant-vowel* (*VCV*) combination using the complete set of Spanish vowels and voiced stop consonants (/*a*/, /*e*/, /*i*/, /*o*/, /*u*/ and /*b*/, /*d*/, /*g*/). Each participant produced the whole corpus on three different sessions taking place on different days (see Methods).

### From continuous dynamics to a discrete motor representation

A visual inspection of the data revealed that the sensors signals remain stable during the utterance of each phoneme and execute rapid excursions during the transitions in order to reach the next state (see Figure 1c); moreover, the signals persist in the same range of values for different vocalization of the same phoneme. This observation invited to hypothesize that each phoneme could be described in a three dimensional discrete space by adjusting thresholds over the signals. This hypothesis was mathematically formalized and tested by using a subset of the data to extract the thresholds and the rest to compute the decoding performance of the phonemic identity.

#### Thresholds

A previous study showed that applying one threshold for each transducer was enough to decode the 5 Spanish vowels^10^. Following the same strategy an extra threshold per signal is added in order to include the stop consonant to the description. Then, the signals were discretized by fitting two thresholds: a vowel threshold *v,* dissociating vowels, and a consonant threshold *c*, dissociating vowels from consonants. A visual exploration of the signals (see Figure 1c for an example, in Supplementary Materials the whole dataset is available) suggested the following rules to fit the thresholds:

Vowels: the lips *v_L_* threshold divides vowels according to the roundness of the lips (above: /*u*/ and /*o*/ below: /*a*/ /*e*/ and /*i*/); the jaw *v_J_* threshold differentiates close (/*u*/ and /*i*/, above) from open (/*a*/ /*e*/ and /*o*/, below) vowels; the tongue *v_T_* threshold separates front vowels (/*e*/ and /*i*/, above) from back (/*a*/ /*o*/ and /*u*/, below) ones.

Consonants: the lips *c_L_* threshold differentiates /*b*/ (above) from all other phonemes (below); the jaw *c_J_* threshold differentiates all stop consonants from all vowels (above: /*g*/ /*b*/ and /d/, below: (/*a*/ /*e*/ /*i*/ /*o*/ and /*u*/); the tongue *c_T_* threshold separates /*d*/ (above) from all other phonemes.

These threshold rules define three regions as the ones shown in Figure 1c in shades of red for the lips, green for the jaw and blue for the tongue. Associating the values 0, 1 and 2 to the transducer signals falling within the light, medium and dark the phonemes could be represented on a discrete 3-dimensional motor space as shown in Figure 1d, were the first coordinate represents the lips state, the second the jaw and the third the tongue. Each vowel is represented by a unique vector: /*a*/=(0,0,0); /*e*/=(0,0,1); /*i*/=(0,1,1); /*o*/=(1,0,0); /*u*/= (1,1,0); while the consonants have multiple representations: /*b*/=(2,*x*,*y*), /*d*/=(*y*,*x*,2) and /*g*/=(*x*,2,*y*), where *x* is 0, 1 or 2, and *y* is 0 or 1.

The thresholds were fixed independently for each transducer signal by choosing the value that better accomplishes the previous described rules (see Methods).

#### Mathematical description for the discretization process

The transformation from continuous transducer signals to discrete values can be mathematically accomplished through saturating functions of the form:

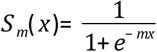

This function goes from zero to one in a small interval around *x*=0, whose size is inversely proportional to *m*. Then, *S*_∞_(*h*(*t*)− *v*) is zero for *h*(*t*)<*v* and one for *h*(*t*)>*v.* These are the conditions that define the binary coordinates for vowels. Using the transducer signals *h_L_*(*t*), *h_J_*(*t*), *h_T_*(*t*) and the threshold values *v_L_, v_J_* and *v_T_* for the lips, jaw and tongue respectively, the vowels read:

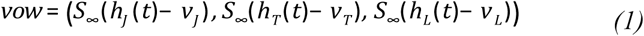

Plosive consonants represent articulatory activations reaching the dark areas of Figure 1c, assigned to the value 2. In order to include them to the description an extra saturating function was added to each coordinate, using the consonant thresholds *c_J_*, *c_T_* and *c_L_*. Following the previous notation, phonemes (either vowels or consonants) can be represented in the discrete space directly from the transducer signals as:

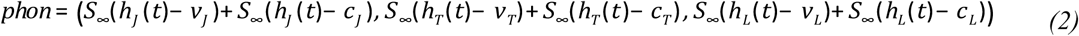

#### Decoding performance: Intra-subject & intra-session

To perform the decoding it is necessary to define the threshold values. In this case, one set of thresholds was adjusted for each participant and session. More precisely, the first 15 *VCV*s of the session were used as training set, i.e. to fix the thresholds (see Table S1 for the numerical values). The following 60 *VCV*s of the corresponding session were used as the test set, i.e. to calculate the decoding performance using the thresholds optimized on the training set.

Figure 2a shows the confusion matrix obtained by averaging the decoding performance across participants and sessions (see Figure S1 for each participant’s confusion matrix). Every phoneme is decoded with performances well above chance levels. This result validates the discretization strategy and discloses a discrete encoding of the phonemic identity in the continuous vocal tract movements.

**Figure 2.**
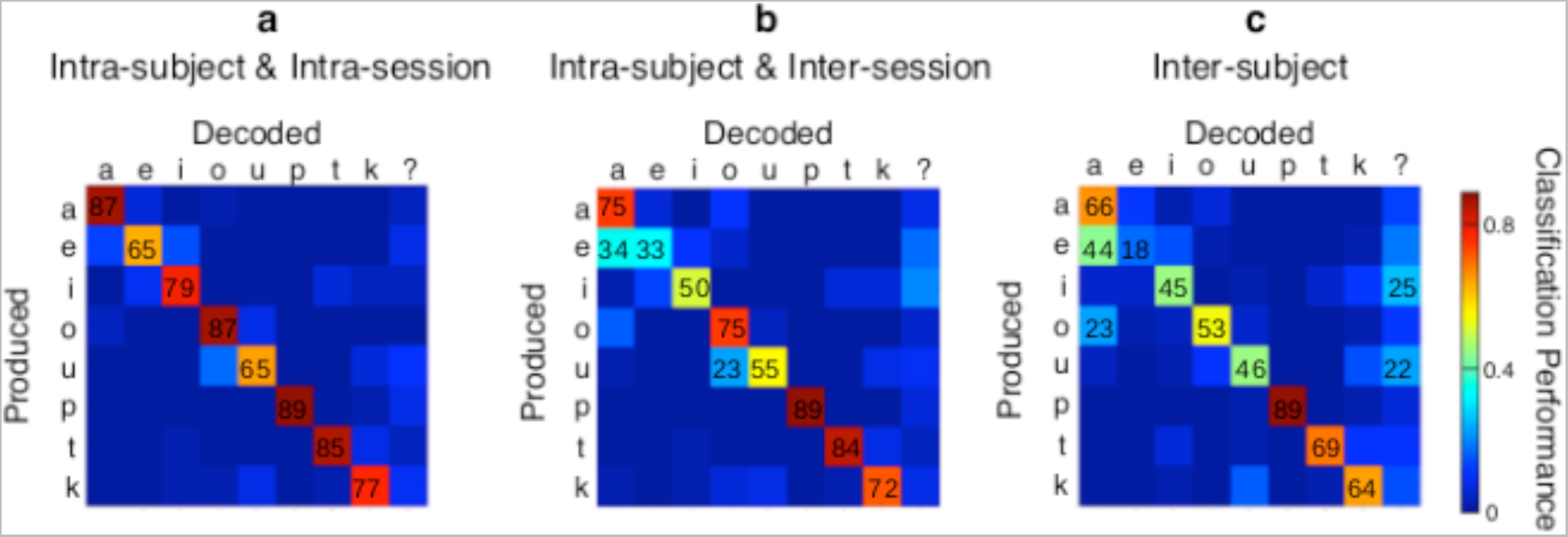
Decoding performance of the discrete representation. A) Confusion matrix using one threshold per session. Average across sessions and participants. B) Confusion matrix using one set of thresholds per participant. Average across participants. C) Confusion matrix computed using one set of thresholds for all participants. Numerical values above 20% are displayed. Thresholds values and matrix means with standard deviations are shown in Table S1 and S2, respectively, at Supplementary Materials.

#### Decoding performance: Intra-subject & inter-session

The previous result leads to the question of whether thresholds can be defined for each participant, *independently* of the variations in the device mounting across sessions. To explore this, the *VCV* data of all sessions were pooled together for each participant. Then the 10% of the data was used to adjust thresholds and tested the performance on the rest of the data. More specifically, a 50-fold cross-validation was performed over each subject’s data set. The confusion matrix of Figure 2b exposes the confusion matrix obtained by averaging the decoding performance across participants (see Figure S2 for individual participant’s confusion matrixes and Table S1 for the mean value and standard deviation of the 50 thresholds). The performance remained well above chance for every phoneme, with the only one exception of the vowel /*e*/, that can be confused with /*a*/. As shown in Figure 1 d, these two vowels are distinguished by the state of the tongue, the articulator for which the mounting of the device is more difficult to standardize.

#### Decoding performance: Inter-subject & inter-session

Next the robustness of the configuration, regardless anatomical differences amongst subjects, was tested. Therefore the *VCV* data from all sessions and participants were pooled together and the 10% of the data, with a 50-fold cross-validation, was used to fix thresholds (see Table S1). The confusion matrix of Figure 2c represent the average values obtained from 50-fold cross-validation. As in the previous case, the vowel /*e*/ was mistaken for /*a*/, revealing that the mounting of the magnet on the tongue needs to be treated with a more fine protocol. This results shows that the discretization strategy is robust even while dealing with different anatomies, suggesting that the encoding of the sounds of language in a low-dimensional discrete motor space represents a general property of the speech production system.

#### Occupation of the consonant’s free states

As pointed out before, vowels and consonants have different *ranks* in the discrete representation: while each vowel is represented by a vertex of the cube of Figure 1d, each consonant is compatible with many states, shown as the points on the 'walls' surrounding the cube. The occupation levels of those states were explored. The discrete state for each consonant was computed using the intra-subject and intra-session decoding, for all participants and sessions, and just the *VCVs* that were correctly decoding were kept for this analysis. The occupations of the different consonantal states are shown in Figure 3a.

**Figure 3.**
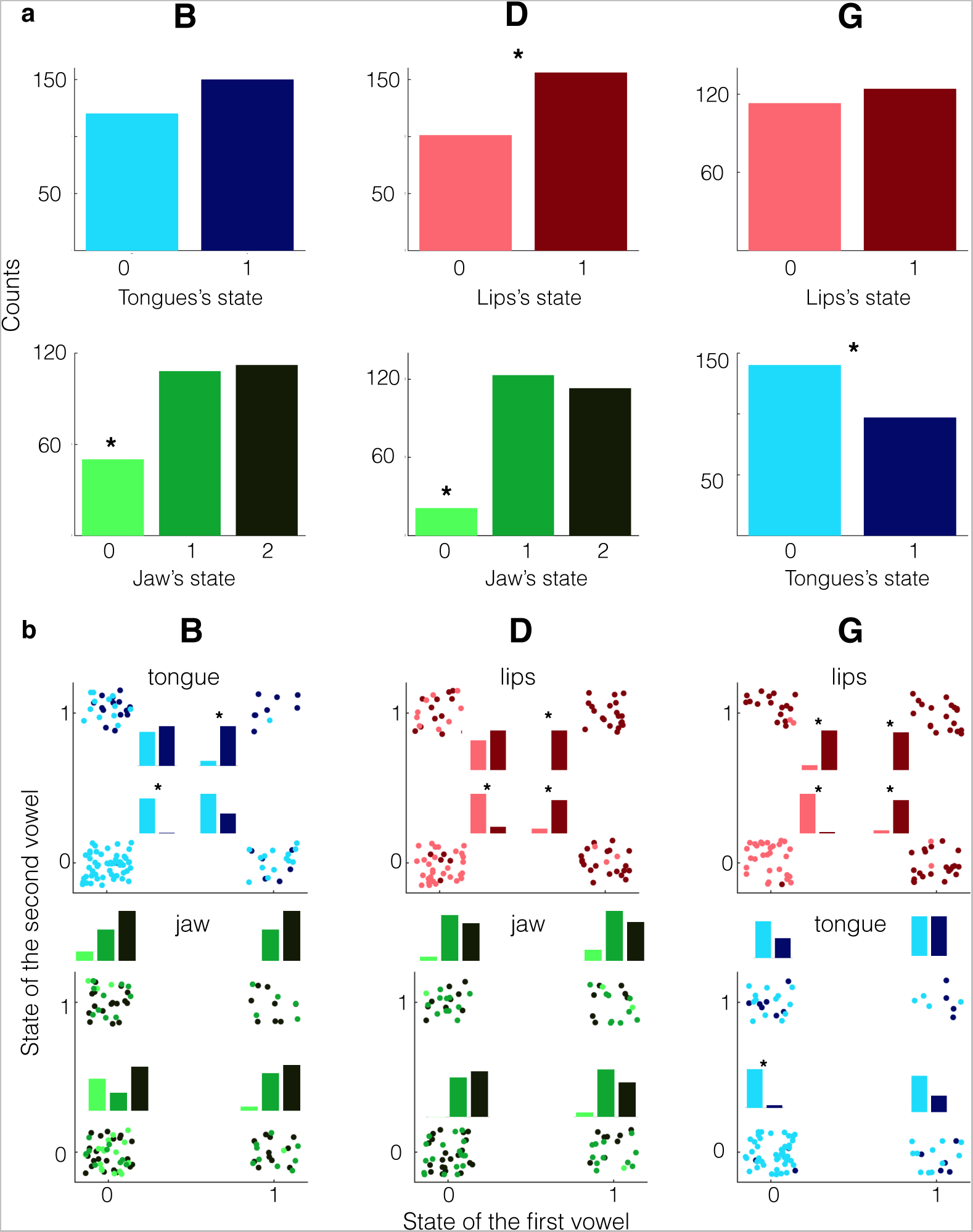
Occupation of the free coordinates. A) Counts of occurrence of each free state, for the complete pool of sessions and participants using the intra-subject and intra-session thresholds for the discretization. Columns represent the different consonants (/*b*/, /*d*/ and /*g*/) and row the free coordinate. Asterisks indicate significant difference between states (p<0.01 binomial test). B) Occupation levels of the free coordinates as a function of the surrounding vowels. The *y*-axis defines the incoming vowel and the *x* the previous one. For each combination of levels the individual vocalizations -dots- and the corresponding histograms are displayed. The color identifies the articulator -red: lips; green: jaw and blue: tongue- and the intensity the given state -light: zero; middle: one and dark: two. Asterisks indicate significant difference between states (p<0.001).

The /*b*/ is defined by the lips in state 2; the tongue and the jaw are free coordinates. The state 2 has not been observed in the tongue, and is presumably incompatible with the motor gesture of this consonant, however no significant differences were found between the states 0 and 1 (binomial test with equal probabilities, p=0.1). Similarly, for the jaw coordinate the state 0 is underrepresented, with an occupation of the 18%, below the chance level of 1/3 (binomial test, p<0.001). The /*d*/ is defined by the tongue in state 2; the lips and the jaw are free coordinates. The lips show a dominance of the state 1 over the 0 (binomial test, p<0.001), and the state 0 of the jaw is significantly less populated than the others with an occupation of the 8%, lower than the chance level of 1/3 (binomial test, p<0.001). The /*g*/ has free lips and tongue coordinates. The lips show no significant differences between the states 0 and 1 (binomial test with equal probabilities, p=0.52), and the state 0 was preferred for the tongue (binomial test with equal probabilities, p=0.006).

A well-known effect in the experimental phonology field is the *coarticulation*; this effect implies that the articulation of the consonants is modified by the neighboring vowels^11^. The occupation levels of the consonants as a function of their surrounding vowels were calculated (Figure 3b) and *coarticulation* effects were revealed. The results show that when the surrounding vowels share some of the consonant’s free states, this state is transferred to the consonant. Specifically, when the previous and following vowels share the lip state its value is inherited by the consonants with free lip’s coordinate, being /*d*/ and /*g*/ (p<0.001 for the four binomial tests). Additionally, /*b*/ inherits the state of the tongue of the surrounding vowels when both share that state (p<0.001 for both binomial tests), and when both vowels share the tongue state 0, it is inherited by the /*g*/ (binomial test, p<0.001). No *coarticulation* is presented by the jaw: for /*b*/ and /*d*/ the jaw is homogeneously occupied by the states 1 and 2, regardless of the states of the surrounding vowels.

### Synthesizing intelligible speech from the anatomical recordings

One of the goals of this study is to produce synthetic speech from the recordings of the upper vocal tract movements by driving an articulatory synthesizer with the signals coming from the sensors. Moreover, considering that the discrete representation successfully decodes the phonemes, the synthesizer will be driven by Equation 2, instead of by the raw signals. Since normal speech arises from continuous changes in the vocal tract, rather than instantaneous passages from one configuration to the other, it is necessary to get smooth transitions from one state to the other. Therefore, *^m^*^= ∞^ in Equation 2 were replaced with finite values *m_1_*, *m_2_* and *m_3_*, for the lip, jaw and tongue coordinate respectively (see Methods). Then, the *smooth version* of Equation 2 was used to drive the articulatory synthesizer described in the Methods section. More precisely, audio files were generated by driving the synthesizer with Equation 2 fed with the transducers' time traces produced by one of the participants while uttering /*oga*/, /*agu*/, /*ogu*/, /*oda*/, /*odi*/, /*ide*/, /*ibu*/, /*iba*/, /*obe*/ (examples available as Supplementary Materials). Details of this process can be found in the Methods section.

Finally, to test the intelligibility of the synthetic speech, the samples were presented to 15 participants, who were instructed to write down a *VCV* structure after listened to each audio file. The confusion matrixes obtained from the transcription are shown in Figure 4 for consonants and vowels. All values are above chance levels (33% for consonants and 20% for vowels).

**Figure 4.**
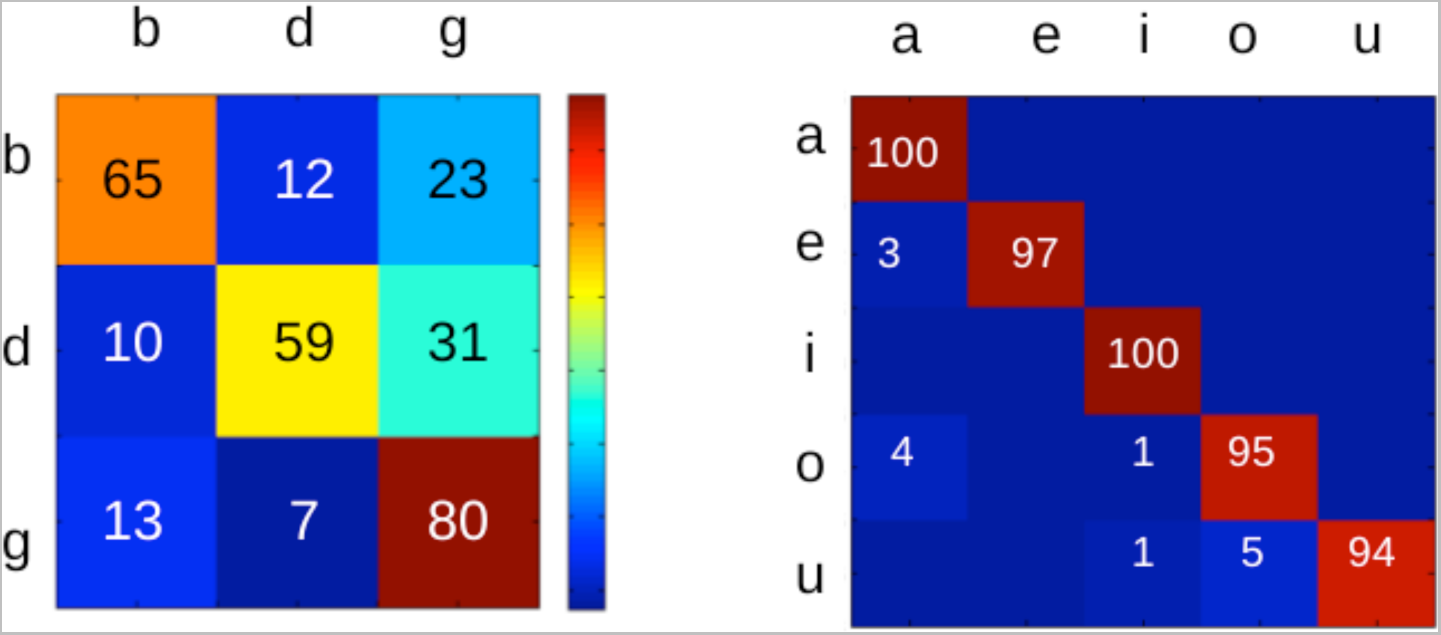
Confusion matrixes for the intelligibility of the synthetic speech. Rows represent the produced phoneme and columns the transcribed one. Right panel: consonants. Left panel: vowels. The color code represents the percentage of responses for each condition.

## CONCLUSIONS

The current work presents two main achievements: *(i)* from a scientific point of view, a discrete representation of the vocal gestures for speech is reconstructed from the direct measurements of the continuous upper vocal tract movements; *(ii)* from a technological point of view, a new device capable of monitoring the upper vocal tract movements during continuous speech is conceived.

### A discrete motor space representation

This study shows that discretizing three continuous signals given by the movements of the main articulators of the vocal tract while vocalizing *VCV* structures is enough to recover the phonemic information. This result represents a follow up of a previous work were the same experimental procedure was applied over isolated vowels. Here, the discrete motor vowel representation is validated during continuous speech and the Spanish stop consonants are integrated to the description.

In order to recover the vowel’s identity just one threshold per signal is needed. Thus, the vowels are represented in the discrete motor space as the corners of a cube. Curiously, the dimension of the *vowel cube* (eight) is in agreement with the number of Cardinal Vowels^12^, a set of vocalic sounds used by the phoneticians to approximate the whole set of cross-language vowels. This dimension match suggests that the discrete motor states captured by this study could represent the basic motor gestures of vowels. Moreover, the state on each *articulator’s transducer* corresponds to an extreme value along the two-dimensional coordinate system used by the International Phonetic Alphabet to describe vowels. Interestingly, the same discrete representation for vowels could be recover from direct measurements of human brain activity during vocalizations^13^.

The consonants chosen for this study were /*b d g*/. They cause a complete occlusion of the vocal tract produced by the constriction gesture of one of the three independent oral articulator sets (lips for /*b*/, tongue tip for /*d*/ and tongue body for /*g*/); and they have been suggested as the basic units of the articulatory gestures^14^. Therefor, they appear as the natural candidates to study the presence of discrete information within the continuous movements of the oral tract.

According to the International Phonetic Alphabet two features define the consonants: the place and the manner of articulation^2^. This work focused in the *place dimension*, since the phoneme selected cover the three main groups used to describe the place of articulation of consonants: labial (/*b*/), coronal (/*d*/) and dorsal (/*g*/). The results revealed the plausibility of recovering the main place of articulation just by monitoring (and discretizing) the kinematics of three points in the upper vocal tract. Similarly, at brain level, spatial patterns of activity during speech show a hierarchal organization of phonemes by articulatory features; where the primary tier of organization is defined by the three major place of articulation: dorsal, labial or coronal^3^.

How the discrete phonemic identity is encoded in the continuous movements producing speech remains an open question for the experimental phonology. The presence of compositional motor units into the continuous articulatory movements has been widely theorized on the literature^15–17^; and some experimental evidence has supported this hypothesis^18–20.^ To our knowledge, this works is among the firsts to recover a discrete representation for the complete set of Spanish vowels and stop consonants from direct measurements of the upper vocal tract movements during continuous speech. This result fills a gap in the comprehension of the speech process by bounding the discrete information at brain level with the discrete phonemic space through the experimental evidence of discrete motor gestures of speech.

### A novel device to track the vocal tract movements during continuous speech

After a short training session the system presented in this work -the electronic device used with the threshold strategy to feed the articulatory synthesizer- produces intelligible speech. While compared with other synthesizers^21,22^ generating speech from vocal movements this one presents the following advantages: *(i)* it is portable; *(ii)* the motor gestures involved in the control of the articulatory model are saturating signals, a shared feature with the brain activation during speech, which represents a clear benefit for brain-computer interface applications.

From a more general point of view, this implementation represents an alternative to the extended strategy used in the bioprosthetic field: large amounts of *non-specific* physiological data processed by statistical algorithms to extract relevant features for vocal instructions^23,24^. Instead, in the current approach a small set of recordings from the movements of the speech articulators, in conjunction with a *threshold strategy,* are used to control a biophysical model of the vocal system.

Although this approach shows potential benefits for bioprothetic applications, further work is needed to optimize the system. On one hand, the mounting protocol for the tongue should tighten up to get stable threshold across sessions. On the other, the protocol should be refined to include the whole consonants data set. Arguably, the manner of articulation could be integrated by including different sets of thresholds; and increasing the number of magnet-transducer sets mounted on the vocal tract could retrieve other places of articulation. Regarding the vowels, the current vocalic space is complete for the Spanish language, and has the same dimension as the cardinal vowels suggesting that it would be enough to produce *intelligible* speech in any language^25^.

The state of the art of the techniques used to monitor the articulatory movements during speech remained stagnant during the last decades, with some exceptions employing different technologies to measure the different articulators^24^. The standard method used to track the articulator’s displacements during speech is the EMA^8,9^. This technique has been prove to provide very accurate recordings ^26-28^ at the expenses of being non portable and expensive. Here, a novel method is introduced and proved to be able to capture the identity of the uttered phoneme, to detect *coarticulation* effects and to correctly drive an articulatory speech synthesizer. This device presents two main advantages: it is portable and is non-expensive. The portability of the system makes it suitable for bioprosthetic applications; and, crucially, because of the low cost of its components, it could significantly improve the speech research done in non-developed countries.

## METHODS

### Ethics Statements

All the participants signed a written consent to participate in the experiments, which were approved by the CEPI ethics committee of Hospital Italiano de Buenos Aires, qualified by ICH (FDA -USA, European Community, Japan) IRb00003580.

### Participants

Four individuals (1 female) within an age range of 29±6 years and no motor or vocal impairments participated in the recordings of anatomical and speech sound data. They were all native Spanish speakers, graduate students working at the University of Buenos Aires. Fifteen participants (9 females) native Spanish speakers participated in the audio tests.

### Experimental device for the anatomical recordings

Participants wore a removable plastic dental cast (1 mm thick) during the experiments (dental casts of the superior and inferior dentures of each participant were supplied by the participants’ dentists). Hall effect transducers (Ratiometric Linear Hall Effect Sensor ICs for High-Temperature Operation, A1323 Allegro) and small biocompatible Sm-Co magnets (1–3g) were mounted in different positions of the upper vocal tract for each participant (see Fig 1a). The dimensions of the transducer are 3x4x1 mm (for height, length and thickness, respectively). The transducer wires (Subminiature Lead Wire TDQ 44, Phoenix Wire Inc.) coated with a plastic tube (Silastic, Laboratory Tubing 0.76 mm61.65 mm) were connected to a variable amplifier (2–30x), low-pass filtered (20 Hz) and recorded with a computer at a 44100Hz sampling rate. Transducer wires and magnets were glued to the plastic replica using cyanoacrylate glue (Crazyglue, Archer, Fort Worth, TX). Denture adhesive (Fixodent Original Denture Adhesive Cream 2.4 Oz) was used to attach magnets to the tongue and medical paper tape (3M Micropore Medical Tape) to fix the transducers to the lips.

Details of the configuration of the 3 magnet-transducer sets shown in figure 1b:

#### Red, lips

One cylindrical magnet (3.0 mm diameter and 1.5 mm height) was glued to the dental cast between the lower central incisors. Another one (5.0 mm diameter and 1.0 mm height) was fixed with medical paper tape at the center of upper lip. The transducer was attached at the center of the lower lip. The magnets were oriented in such a way that its magnetic field has opposite signs in the privileged axis of the transducer.

#### Green, jaw

A spherical magnet (5.0 mm diameter) and the transducer were glued to the dental casts, in the space between the canine and the first premolar of the upper and lower teeth respectively.

#### Blue, tongue

A cylindrical magnet (5.0 mm diameter and 1.0 mm height) was attached at a distance of about 15 mm from the tip of the tongue, using a small amount of denture adhesive. The transducer was glued to the dental plastic replica, at the hard palate, approximately 10 mm right over the superior teeth (sagittal plane). Transducer wire was glued to the plastic replica and routed away to allow free mouth movements.

### Recording sessions

Each participant completed 3 sessions on different days during a period of one month. Participants with the device mounted, sat in a silent room 20 cm away from a microphone and in front a computer screen. They were instructed to repeat the different VCVs structures that prompted on the screen starting from (and ending to) a comfortable closed mouth configuration. Each session incorporates the production of 75 VCVs strings, including the whole set of Spanish vowels (/*a*/, /*e*/, /*i*/, /*o*/ and /*u*/) and voiced plosive consonants (/*b*/, /*d*/ and /*g*/). The first 15 VCVs were defined as the training set and include the same number of each phoneme (5 /*b*/, 5/*g*/, 5 /*d*/, 6 samples of each vowel). The same design was applied for the next 60 vocalizations (20 samples of each consonant, 24 for each vowel). The VCVs order was randomized for each session. The speech sounds were recorded simultaneously with the three signals of the transducers.

The consonants chose for this study are produced by the activation of different articulators and represents the main hierarchies used by IPA^14^ to categorize the consonants according to its place of articulation: /*b*/ labial, /*d*/ coronal and dorsal /*g*/.”

### Preprocessing and thresholds fitting

Each transducer’s signal was subsampled at 441 Hz and preprocessed within each session data set. The preprocessing consisted in subtracting the resting state and dividing by the maximum absolute value. The resting state value was estimated as the average of the 50 ms previous and posterior of each vocalization. This process sets the signal’s value between −1 and 1.

Each threshold was fixed independently, by choosing the value maximizing the decoding performance over the training data set, according to the rules detailed above in the *Thresholds* section. For example, for the tongue vocalic threshold the rule is: /*e*/ and /*i*/ above (1) and /*a*/ /*o*/ and /*u*/ below (0). The [-1;1] range was discretized in 200 evenly spaced values, each value was tested as threshold over the tongue’s signals of the complete testing data set. The value with the highest performance sorting the data according to the rule was selected as threshold.

### Articulatory synthesizer

During the production of voiced sounds, the vocal folds oscillate producing a stereotyped airflow waveform^29^ that can be approximated by relaxation oscillations^30^ such as the produced by a van der Pol system:

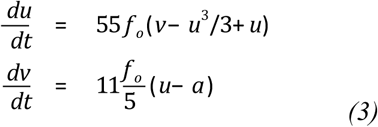

The glottal airflow is the variable *u* for *u*>0, and *u*=0 else. The fundamental frequency of the glottal flow is *f*_0_ (Hz) and the oscillations' onset is attained for *a*>-1.

The pressure perturbations produced by the injection of airflow at the entrance of the tract propagate along the vocal tract. The propagation of sound's waves in a pipe of variable cross section *A*(*x*) follows a partial differential equation^31^. Approximations have been proposed to replace this equation by a series of coupled ordinary differential equations, as the wave-reflection model^32–34^ and the transmission line analog^35^. Those models approximate the pipe as a concatenation of *N*=44 tubes of fixed cross-section *A_i_* and length *l_i_*. In the transmission line analog, the sound propagation along each tube follows the same equations as the circuit shown in Figure 5, where the current plays the role of the airflow *u* and the voltage the role of sound pressure *p.* The flows *u*_1_, *u*_2_ and *u*_3_ along the meshes displayed in Figure 5, follow the equations:

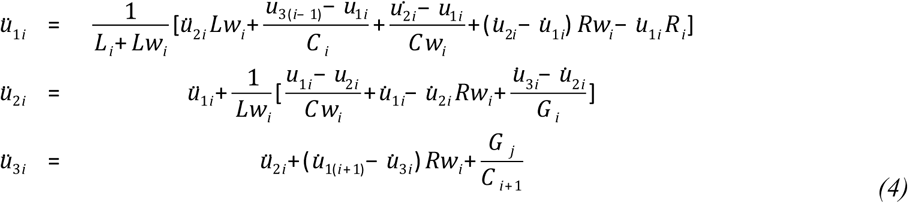

**Figure 5.**
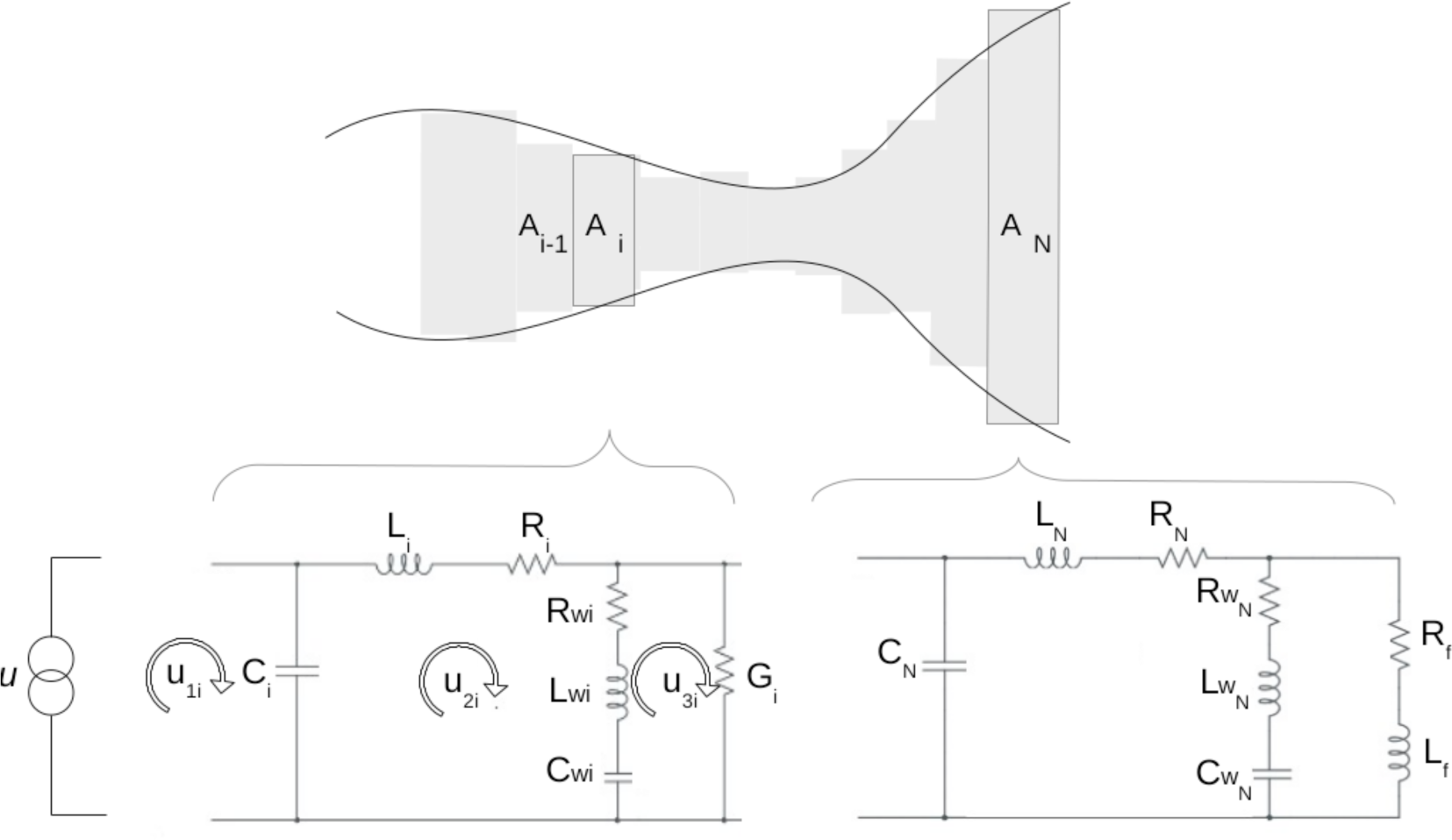
Articulatory synthesizer. A) Sketch of the model used for the vocal system. The glottal airflow (*u*) is obtained by solving Van del Pol system^30^. The vocal tract is approximated by a concatenation of *N*=44 tubes of different cross-sectional areas *A_i_* and lengths *l_i_*. The lower panels show the equivalent electrical transmission line used to simulate the propagation of the sound along the *ith* and the last tube (*N*), respectively.

The electric components *L_i_*, *C_i_*, *R_i_* and *G_i_* are functions of the cross-section and length of the small *i-*th tube and represent the acoustic inertance, the air compressibility, the power dissipated in viscous friction and the heat conduction at the tube wall, respectively. The components marked with *w* account for the vibration of the walls of the tract and the mesh for the last tube (*i=N)* includes elements (*R_f_* and *L_f_*) that account for the mouth radiation. For a complete description of the model and the numerical parameters see Flanagan et al., 2008^35^.

Speech was synthesized using the glottal airflow *u* as input for the first circuit *i*=1 (Figure 5), which represents the entrance to the tract. Then, the propagation of the sound along the vocal tract was modeled through *N* sets of Equation 4.

### Discrete states to vocal tract anatomies

The shape of the vocal tract can be mathematically described by its cross-sectional area *A*(*x*) at distance *x* from the glottal exit to the mouth. Moreover, previous works^34,36,37^ developed a representation in which the vocal tract shape *A*(*x*) for any vowel and plosive consonant can be expressed as

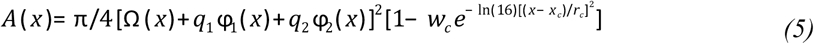

The first factor in square brackets represents the shape of the vocal tract for vowels, the *vowel substrate*. The function *Ω(x)* is the neutral vocal tract, and the functions *ϕ*_1_(*x*) and *ϕ*_2_(*x*) are the first empirical modes of an orthogonal decomposition calculated over a corpus of MRI anatomical data for vowels^37^. The shape of the vowels is determined by fixing just two coefficients, *q*_1_ and *q*_2_^36^. The second factor represents the consonant occlusion. It takes the value 1 except for an interval of width *r*_c_ around the point *x*=*x*_c_, where it smoothly decreases to 1-*w*_c_ and therefore represents a local constriction in the vocal tract, active during a plosive consonant. Basically, *w_c_* goes to one during the consonant occlusion and stays in zero in other case.

This description of the anatomy of the vocal tract fits well with our discrete representation. A previous study^10^ showed that a simple map connects the discrete space and the morphology of the vocal tract for vowels. It is carried out by a simple affine transformation defined by:

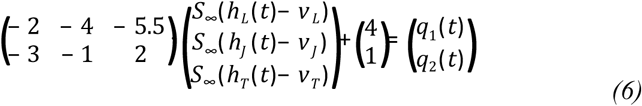

The numerical values of the transformation were phenomenologically found to correctly map the discrete states to the vowel coefficients *q*_1_ and *q*_2_. Together, Equations 5 and 6 allow the reconstruction of the vocal tract shape of the different vowels from the transducer signals.

During plosive consonants, the vocal tract is occluded at different locations. In our description, this corresponds to have a value 2 in one or more coordinates, which means that the transducers signal cross the consonant threshold *c*. The saturating functions with the consonant threshold were used to control the parameter *w*_c_ of Equation 5 that controls the constriction. More specifically, the following Equations were used to generate the consonants:

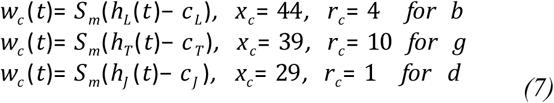

The values of *x*_c_ and *r*_c_ are in units of a vocal tract segmented in 44 parts, starting from the vocal tract entrance (*x*_c_= 1) to the mouth (*x*_c_=44).

This completes the path that goes from discretized transducer signals *h_J_*(*t*), *h_T_*(*t*) and *h_L_*(*t*) to the shape of the vocal tract *A*(*x*,*t*) for vowels and plosive consonants.

### Vocal tract dynamics driven by transducers' data

To produce continuous changes in a virtual vocal tract controlled by the transducers, it is necessary to replace the infinitely step functions in Equation 6 and 7 by smooth transitions from 0 to 1. Therefor, the condition *m*=∞ is replaced by finite steepness values *m*_1_, *m*_2_ and *m*_3_. The values used to synthesize continuous speech were *m*_1_=300, *m*_2_=300 and *m*_3_=900 for lips, tongue and jaw, respectively. These numerical values were manually fixed with the following constrain: applying Equation 6 over the recorded signals during the stable part of the vowels, and using the obtained (*q_1_, q_2_*) to synthesize speech should produce recognizable vowels. This process is explained below.

First, the mean values of the transducer signals during the production of vowels for one participant were computed (left panel Figure 6). More precisely, just the set of corrected decoded vowels for subject 1, using the intersession threshold, were selected. Second, different exploratory sets of (*m*_1_, *m*_2_,*m*_3_) were used to calculate the corresponding (*q_1_*, *q_2_*), by means of Equation 6. Then, the given vocal tract shapes (*A(x)* in Equation 5) could be reconstructed and the vocalic sounds were synthetized, from which the first two formants were extracted using Praat^38^. Each sets of (*m*_1_,*m*_2_,*m*_3_) produce a different map going from the sensor space to the formants space (Figure 6)

**Figure 6.**
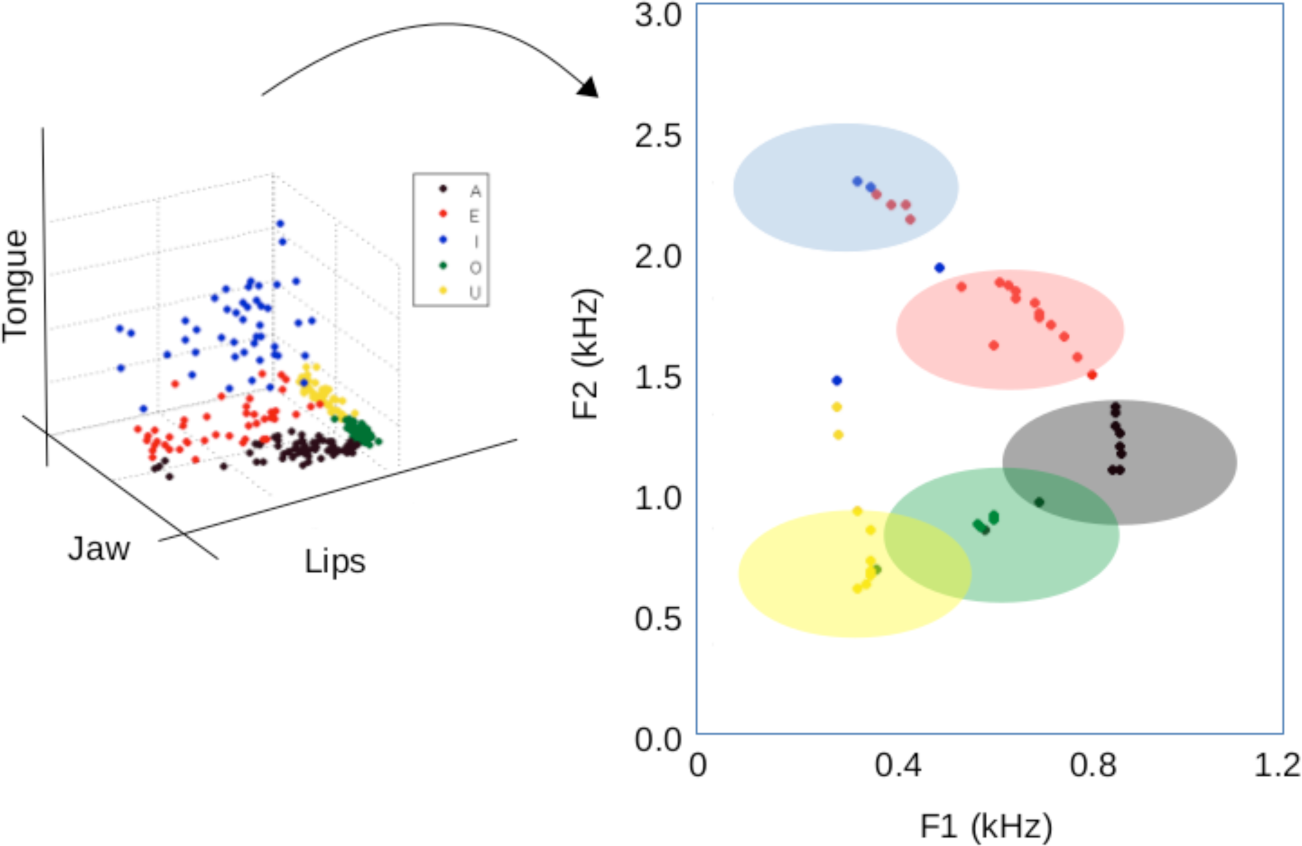
Mapping the transducer signals into the formants space. The left panel shows the mean values *h_J_*, *h_T_* and *h_L_* of the transducer signals during the utterance of vowels by one participant. Each dot represents a vowel, colors stands for the vowel’s identity (/*a*/ is black, /*e*/ red, /*i*/ blue, /*o*/ green, /*u*/ yellow). The right panel maps each point in the transducers space to the formants space as described above, for (*m*_1_, *m*_2_,*m*_3_)=(400,400,400). The shaded regions represent the experimental variation in the formants *F*_1_ and *F*_2_ for Spanish vowels^39^.

The first two formants of a vocalic sound defines its identity^29^; its variability for real vocalizations of Spanish vowels is represented by the shaded areas on the right panel of Figure 6 according to previous reported results^39^. The chose steepness values (*m*_1_=300, *m*_2_=300 and *m*_3_=900) map more than 90% of the transducer data into the experimental (*F_1_,F_2_*) regions.

### Synthetic speech

To synthesize speech it is necessary to solve the set of equations described in the *Articulatory synthesizer* section (Equations 3 and 4). Moreover, for simulating the glottal flow a value for the pitch (*f_0_*) is needed; and to solve the electric analog for the discretized tube the vocal tract shape. The pitch value was extracted using Praat^38^ from the speech recordings. And by means of Equation 5, 6 and 7 it is possible to reconstruct the time evolution of the vocal tract shape from the recording signals. Videos of the vocal tract dynamics driven by the transducers are available at Supplementary Materials for different VCV structures.

Audio wav files were generated using the transducers' time traces and the pitch contours produced by one of the participants while vocalizing: /*oga*/, /*agu*/, /*ogu*/, /*oda*/, /*odi*/, /*ide*/, /*ibu*/, /*iba*/, /*obe*/ (data available as Supplementary Materials). The audio files were generated by integration of Equations 3 and 4 in *N*=44 tubes, using a Runge-Kutta 4 algorithm^40^ coded in C at a sampling rate of 44.1 kHz. The sound intensity of the files was equalized at 50 dB.

Fifteen participants using headphones (Sennheister HD202) listened to the synthetic speech trials in random order. They were instructed to write down a *VCV* structure after listened to each audio file. The experiment was written in Psychtoolbox^41^.

## Aknowledgements

This work describes research partially funded by CONICET, ANCyT, UBA.

## Author contributions

GM conceived the experiments; GM, MFA and MAT designed the experiments; MFA ran the experiments; DRB, MFA and MT coded the synthesizer; MFA and MT analyzed the data and wrote the manuscript.

